# gorder: Comprehensive tool for calculating lipid order parameters from molecular simulations

**DOI:** 10.1101/2025.05.07.652627

**Authors:** Ladislav Bartoš, Peter Pajtinka, Robert Vácha

## Abstract

Lipid order parameters are an important metric for quantifying the molecular structure of biological membranes. They can be derived from both molecular simulations and experimental measurements, enabling robust comparisons between the two. Although methods for calculating lipid order parameters from molecular dynamics simulations of membrane systems at various resolutions are well established, a comprehensive and user-friendly package for these calculations is lacking, which has even led some researchers to use tools that are known to perform the calculations incorrectly. To address this, we have developed `gorder`, an analysis tool capable of calculating lipid order parameters in atomistic, united-atom, and coarse-grained systems, compatible with any force field, and applicable to both planar and curved membrane geometries. `gorder` is designed to be fast and versatile, providing a unified solution for lipid order calculations. The tool is freely available from crates.io/crates/gorder and github.com/Ladme/gorder under the MIT License.

## Metadata

**Table 1:** Code metadata.

| Nr. | Code metadata description | Metadata |
| --- | --- | --- |
| C1 | Current code version | v1.0.0 |
| C2 | Permanent link to code/repository used for this code version | https://github.com/Ladme/gorder |
| C3 | Permanent link to Reproducible Capsule | – |
| C4 | Legal Code License | MIT License |
| C5 | Code versioning system used | git |
| C6 | Software code languages, tools, and services used | Rust, C, Python |
| C7 | Compilation requirements, operating environments & dependencies | required: rustc, cargo, gcc/-clang/msvc; recommended: python, uv |
| C8 | If available, link to developer documentation/manual | https://ladme.github.io/gorder-manual/ |
| C9 | Support email for questions |  |

## 1. Motivation and significance

Lipid order parameters quantify the degree of ordering in phospholipid membranes. In the field of molecular dynamics, they are widely used for assessing force field accuracy [1, 2, 3, 4, 5, 6, 7], studying membrane structures [8, 9, 10, 11, 12, 13, 14, 15], tracking phase transitions [16, 17], and various other applications [18, 19, 20]. The term “lipid order parameters” typically refers to one of two related but distinct metrics: *S_CH_* and *S_CC_*order parameters.

The *S_CH_* (sometimes referred to as *S_CD_*) order parameters are associated with bonds between heavy atoms (typically carbons) and hydrogens. The absolute values of *S_CH_* order parameters can also be obtained experimentally using quadrupolar splitting from deuterium NMR experiments [21, 22] or dipolar splitting measured using carbon-13 NMR experiments [23, 24, 25], rendering this metric particularly useful.

For coarse-grained systems, where multiple heavy atoms and their bonded hydrogens are coalesced into a single bead, this metric is not applicable. Instead, the *S_CC_*order parameters can be used, which relate to bonds between two consecutive beads of the lipid molecule [26]. While *S_CC_* order parameters are not directly quantitatively comparable to the experimentally measured *S_CH_* order parameters, they represent the same physical properties.

In the following text, we will refer to *S_CH_* order parameters calculated using explicit hydrogen positions as “atomistic order parameters,” *S_CH_* order parameters calculated using predicted hydrogen positions as “united-atom order parameters,” and *S_CC_* order parameters as “coarse-grained order parameters.”

The calculation of atomistic, united-atom, and coarse-grained order parameters is relatively well established, with various tools available for computing them from molecular dynamics simulations, including those designed for the popular GROMACS simulation package [27].

In the GROMACS community, a widely used tool for calculating *S_CH_* order parameters is `gmx order`, which is included in the GROMACS simulation package. `gmx order` computes *united-atom* order parameters, implicitly predicting hydrogen positions during the calculation, even if the hydrogens are explicitly available. However, it has been demonstrated that `gmx order` does not correctly calculate order parameters for unsaturated carbons [28]. These findings prompted the GROMACS developers to remove this option entirely from the tool, and `gmx order` is now explicitly marketed as a tool applicable exclusively for analyzing fully saturated united-atom lipid tails [29].

Despite this requirement, the tool is frequently employed in scenarios that contradict it. Since 2020 alone, we have identified at least 16 scientific publications where `gmx order` was applied to unsaturated carbons in atomistic systems [30, 31, 32, 33, 34, 35, 36, 37, 38, 39, 40, 41, 42, 43, 44, 45], 3 articles where it was used to calculate order parameters for unsaturated carbons in united-atom systems [46, 47, 48], and even 2 articles where it was used for coarse-grained systems [49, 50], despite the inherent contradiction of predicting hydrogen positions for coarse-grained lipids. Additionally, gmx order is very frequently employed to calculate order parameters for saturated carbons in *atomistic* systems (see, e.g., [51, 52, 53, 54, 55, 56, 57, 58, 59, 60]). While for these saturated carbons, the errors introduced by predicting hydrogen positions are typically small and often negligible, this usage of gmx order still goes against the recommendation.

There are some alternatives to gmx order that are generally better suited for calculating order parameters. For atomistic systems, one can use tools such as scripts provided by NMR Lipids [61], VMD’s `calc op.tcl` script [62], or the `order params` tool included in the LOOS library [63, 64]. For united-atom systems, the `g lomepro` package (available from github.com/vgapsys/g lomepro) or the `buildH` tool [65] can be employed, both of which handle unsaturated carbons correctly. For coarse-grained systems, options include the do-order script (available from cgmartini.nl/docs/downloads/tools/other-tools.html) provided by the developers of the Martini force field [66, 67], the lipyphilic library [68], or the `order` program [69], a tool previously developed by one of the authors of this article. Additionally, some researchers develop in-house tools tailored to their specific needs.

Despite the availability of these alternatives, the use of `gmx order` persists. This is largely due to its direct inclusion in the GROMACS simulation package, making it immediately accessible for many researchers. Other contributing factors may include the challenging installation process of many alternative tools, the significantly faster computation speed of `gmx order` compared to most alternatives, and its perceived force field independence.

Here, we introduce `gorder`, a tool that integrates and enhances the features of existing tools for computing order parameters. `gorder` calculates lipid order parameters for atomistic, united-atom, and coarse-grained systems, independent of force field. Additionally, it offers advanced features such as leaflet-specific order parameter calculations, construction of order parameter maps, or support for membranes with curved geometries. gorder can be easily installed via Rust’s `cargo` package manager, configured using YAML-formatted files, and is—to the best of our knowledge—the fastest tool available for this purpose.

## 2. Software description

### 2.1. Theoretical background

If explicit hydrogen positions are available, *S_CH_* order parameters can be calculated as:

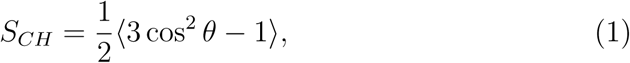

where *θ* is the angle between the C–H bond vector and the bilayer normal, and the angular brackets represent molecular and temporal ensemble averages [70, 71]. The resulting order parameters are typically reported as −*S_CH_*, a convention arising from NMR measurements and followed by gorder as well. Higher values of −*S_CH_*then correspond to more ordered membrane structures.

If hydrogens are not available, their positions need to be predicted. This prediction can be performed either explicitly, where the positions of the hydrogens are first predicted and then the *S_CH_*order parameters are calculated as described in Equation 1, or implicitly, where the hydrogen predictions and the order parameter calculations are performed in a single step. Since gorder uses explicit hydrogen predictions, we do not describe the implicit method here and instead refer the interested reader to [28]. `gorder` predicts hydrogen positions in the same way as the `buildH` tool [65], see the Supporting Information for details.

The calculation of *S_CC_*order parameters also uses Equation 1, but in this case, *θ* represents the angle between the bond connecting two consecutive lipid beads and the bilayer normal [26]. *S_CC_* order parameters are typically reported directly as *S_CC_*without sign inversion, as they naturally follow the same trend as −*S_CH_* .

### 2.2. Software architecture

*Notable dependencies.* The `gorder` tool is built on top of `groan rs` (available from github.com/Ladme/groan rs), an in-house developed Rust library for analyzing molecular dynamics simulations. Several dependencies used by the `groan rs` library are worth noting. TPR file parsing is handled by `minitpr` (available from github.com/Ladme/minitpr), a fully standalone TPR-file parser implemented in pure Rust and developed in-house. XTC file reading is performed by molly (available from git.sr.ht/∼ma3ke/molly), a library also written in Rust, which is reportedly faster than the widely used `xdrfile` library. The `molly` library supports partial XTC frame reading, enabling the program to only read specific atom selections (such as the membrane without the surrounding water), which further enhances analysis speed. However, `groan rs` still relies on `xdrfile` for parsing TRR files.

*Architecture.* The gorder tool uses a three-layered architecture with the input layer handling user input, the logic layer performing the analysis, and the presentation layer managing output of the results. The architecture of `gorder` with its high-level structures is depicted in Figure 1.

**Figure 1:**
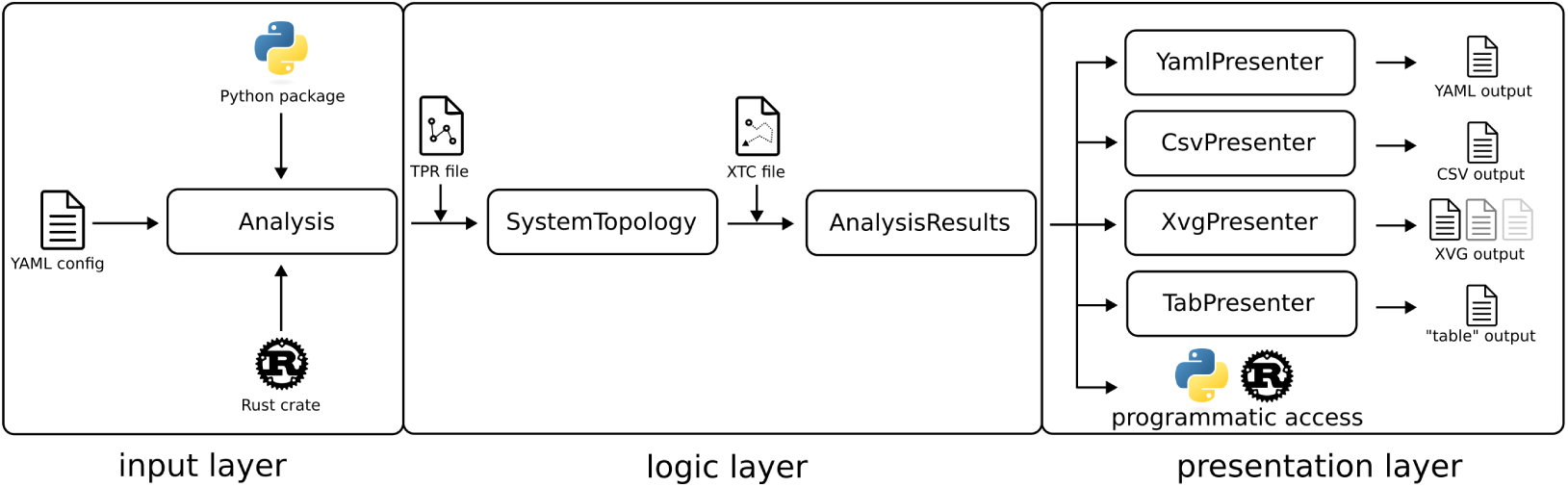
Simplified depiction of the architecture of gorder. In the input layer, the Analysis structure is constructed by either reading a YAML configuration file or using Python/Rust API. In the logic layer, the TPR file (or other supported topology-containing file) is read to construct a `SystemTopology` structure. Subsequently, an XTC file (or other supported trajectory file) is analyzed, generating an AnalysisResults structure. This structure can either be accessed programmatically via Python/Rust API or output in YAML, CSV, XVG, or a custom “table” format using presenter structures (`YamlPresenter, CsvPresenter, XvgPresenter, TabPresenter`).

The analysis itself consists of two main stages. In the first stage, lipid molecules are identified and classified. A collection of atoms connected by bonds is considered to form a single molecule. Molecules are classified as the same type when they contain identical sequences of atoms (with matching ordering in the input topology, matching atom names and matching residue names) and identical sets of bonds (bonded atom pairs). A single molecule may consist of multiple residues, while distinct molecule types can share residue names, if they differ in topology. Individual bonds are categorized into bond types, with order parameters subsequently calculated for each bond type involving atoms selected by the user.

In the second stage, the program processes the selected trajectory frames. For each frame, it iterates through identified molecule types, individual bond types within these molecule types, and specific bond instances, calculating and collecting order parameters. Additional operations may be performed at the level of individual bonds (e.g., selection of bonds in a specific part of the membrane or projecting order parameters onto two-dimensional maps), at the level of individual molecules (e.g., leaflet classification or calculation of dynamic membrane normals), or at the level of the entire system (e.g., leaflet classification using spectral clustering).

Additional details about `gorder’s` internal implementation are provided in the Supporting Information, including descriptions of the algorithms used for membrane leaflet classification and dynamic membrane normal calculations, as well as the multithreading scheme employed by `gorder`.

### 2.3. Software functionalities

`gorder` is able to calculate atomistic, united-atom, and coarse-grained lipid order parameters. It is completely force field independent, which, however, requires the user to provide system connectivity and manually select atoms for the analysis. System connectivity can be supplied by providing a GROMACS TPR file, a PDB file with a connectivity section, or a custom “bonds” file. To simplify atom selection, we implemented a flexible selection language similar to that used by VMD [62], with added support for regular expressions and GROMACS NDX file groups (see the guide at ladme.github.io/gslguide). While `gorder` primarily expects the trajectory to be provided in the XTC file format, TRR and GRO trajectories are also supported. The main functionalities of `gorder` include:

- Reporting order parameters for individual atom types (in atomistic and united-atom systems) as well as for individual bond types.
- Automatic classification of lipid types and their simultaneous analysis, with results reported separately for each lipid type.
- Leaflet-wise order parameter calculations, with automatic or manual assignment of lipids into leaflets.
- Estimating errors using block averaging and reporting on simulation convergence.
- Projecting lipid order parameters onto two-dimensional maps.
- Calculating order parameters for dynamically selected membrane regions.
- Dynamically computing membrane normals based on the current membrane shape, enabling reliable order parameter calculations even for curved membranes, such as vesicles, tubes, or buckled membranes.
- Automatic handling of periodic boundary conditions in orthogonal simulation boxes.
- Multithreading, with binary reproducible output independent of the number of threads used (when run under the same hardware and with the same compiler environment).
- Quality-of-life features, including trajectory concatenation and selection of time ranges for analysis.
- Outputting the results in YAML, CSV, or XVG formats, or in a custom “table” format designed for human readability.

All functionalities of `gorder` are configurable via a YAML-formatted configuration file. `gorder` is available as a command-line application, a Python package, and a Rust crate, facilitating seamless integration into user workflows.

For more information about the functionalities and limitations of `gorder`, we provide a comprehensive manual for the tool, accessible at ladme.github.io/gordermanual or from 10.5281/zenodo.15282375.

## 3. Illustrative examples

This section demonstrates several applications of `gorder` for calculating lipid order parameters across different biological systems. Additional examples can be found in the Supporting Information. For each analysis shown, the `gorder` input parameters are provided in their respective figures. Ready-touse versions of these configurations are available in the Supporting Information listings.

### 3.1. Single-component atomistic membrane

In the first example, we calculated order parameters for both tails of POPC lipids in a planar POPC bilayer. The system contained 256 lipids represented using the CHARMM36 force field [72] and simulated for 200 ns. The trajectory comprised 10,000 frames. The full system consisted of around 64,500 atoms. Complete simulation parameters are provided in the Supporting Information.

We focused on comparing results obtained by `gorder` with those calculated by other software and benchmarking performance, as detailed in Figure 2. Note that while we display only order parameters for carbons here, `gorder` also calculates order parameters for individual carbon-hydrogen bonds, as shown in Figure S2.

**Figure 2:**
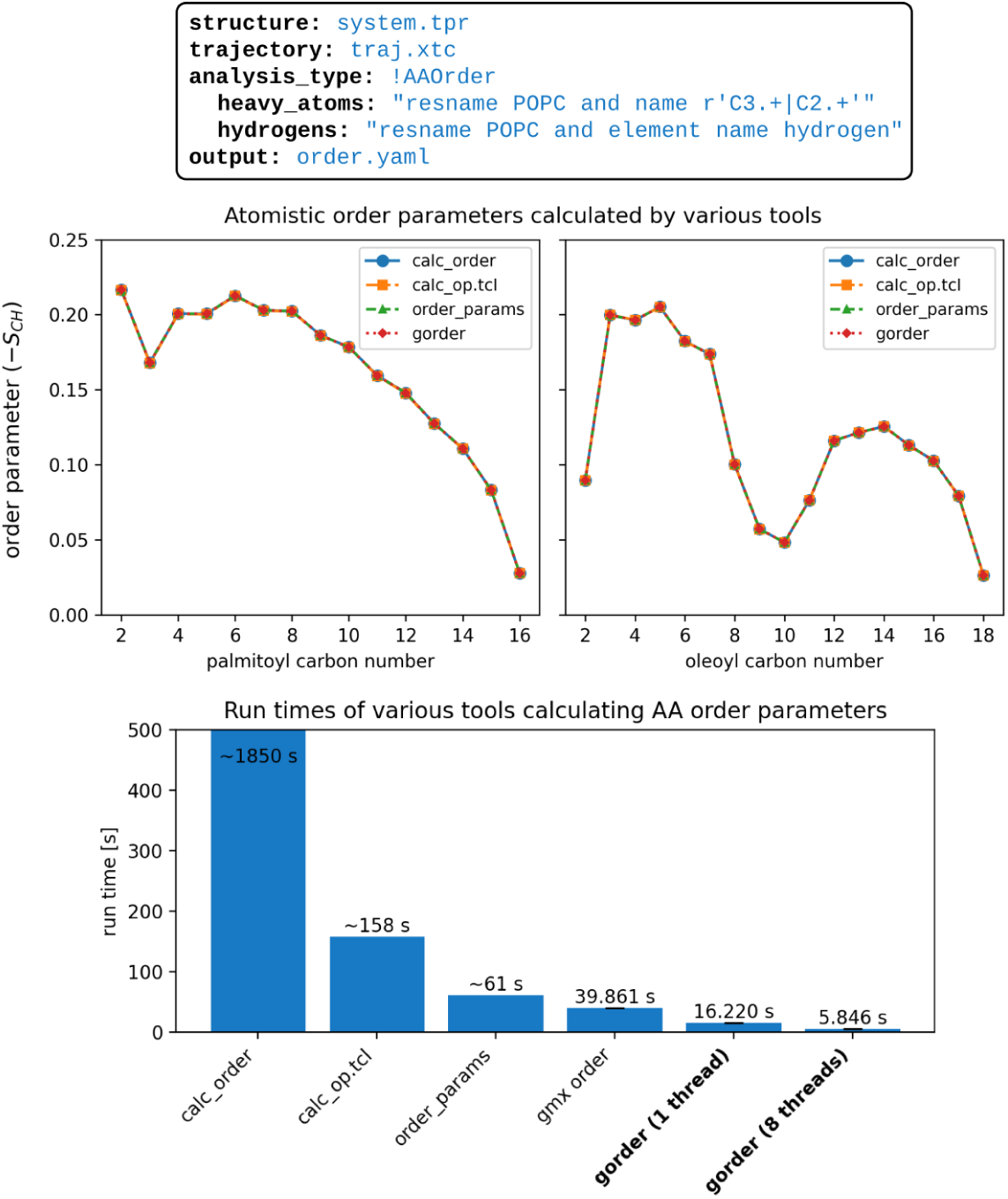
Calculation of order parameters in an atomistic system using `gorder` and comparison with other software. **Top:** Configuration YAML file specifying the analysis parameters for `gorder`. Structure and topology will be read from system.tpr and the trajectory from `traj.xtc`. Atomistic order parameters will be calculated (!AAOrder). The query for heavy atoms selects carbons of palmitoyl and oleoyl tails, while the query for hydrogens selects all hydrogens of POPC molecules. The tool will automatically identify bonds between the selected atoms. Output will be written into order.yaml in YAML format. **Center:** Order parameters calculated by gorder for palmitoyl and oleoyl tails of POPC (red line), compared with results from other tools (NMR Lipids’ calc order script, VMD’s `calc op.tcl`, and LOOS library’s order params). All tools yield consistent results. **Bottom:** Execution times for order parameter calculations of POPC tail carbons across different tools. `gmx order` is included despite being designed only for fully saturated united-atom lipid tails, as it remains commonly used for atomistic systems. Benchmarking was performed on GNU/Linux Mint 20.2 (8-core Intel Core i7-11700, Samsung 870 EVO SSD) with cold cache using hyperfine. `gorder` was compiled using rustc v1.87.0 and gcc v9.4.0. `gorder` and `gmx order` were executed 5 times each, while the other, slower tools were run once. Note: `order params` and `gmx order` require two runs to analyze both lipid tails. Even for a single-tail analysis (single run), `gorder` maintains significantly higher performance with comparable resources (`gorder`: 10.6 s, `gmx order`: 20.7 s).

gorder yields results consistent with other tested tools while being significantly faster. For more complex lipid compositions than presented here, gorder will be even more efficient since it can calculate order parameters separately for all lipid types in one trajectory pass, while all other programs require at least one pass *per* analyzed lipid type. Note that the reported performance values are specific to the hardware and compiler environment detailed in the caption of Figure 2.

We also performed the same analyses and benchmarks for single-component united-atom and coarse-grained phospholipid membranes. In all cases, gorder matches the results of other (correctly performing) analysis tools while maintaining faster performance (see Figures S3 and S4). To further validate hydrogen position predictions in united-atom systems, we also computed united-atom order parameters for an atomistic system and compared them with atomistic order parameters (see Figure S5).

### 3.2. Membrane with complex composition

This example demonstrates gorder’s capability to analyze membranes with complex lipid compositions and estimate calculation errors. We simulated a symmetric CHARMM36 [72] membrane containing eight lipid species: POPC, POPE, POPG, POPA, DOPC, DOPE, DPPC, and PVCL2 (a cardiolipin with two palmitoyl and two oleoyl tails and a 2− charge), with 32 molecules of each type. The analysis was performed using 25,000 frames from a 500 ns trajectory segment. Complete simulation parameters are provided in the Supporting Information.

See Figure 3 for the configuration YAML file specifying analysis parameters and calculated order parameters for individual carbon atoms in each lipid tail. Note that the analysis for all lipid types is performed simultaneously at almost the same total speed as for a single-component membrane (see Figure S6). For evaluation of gorder’s performance when analyzing large lipid molecules (such as cardiolipin), see Figure S7.

**Figure 3:**
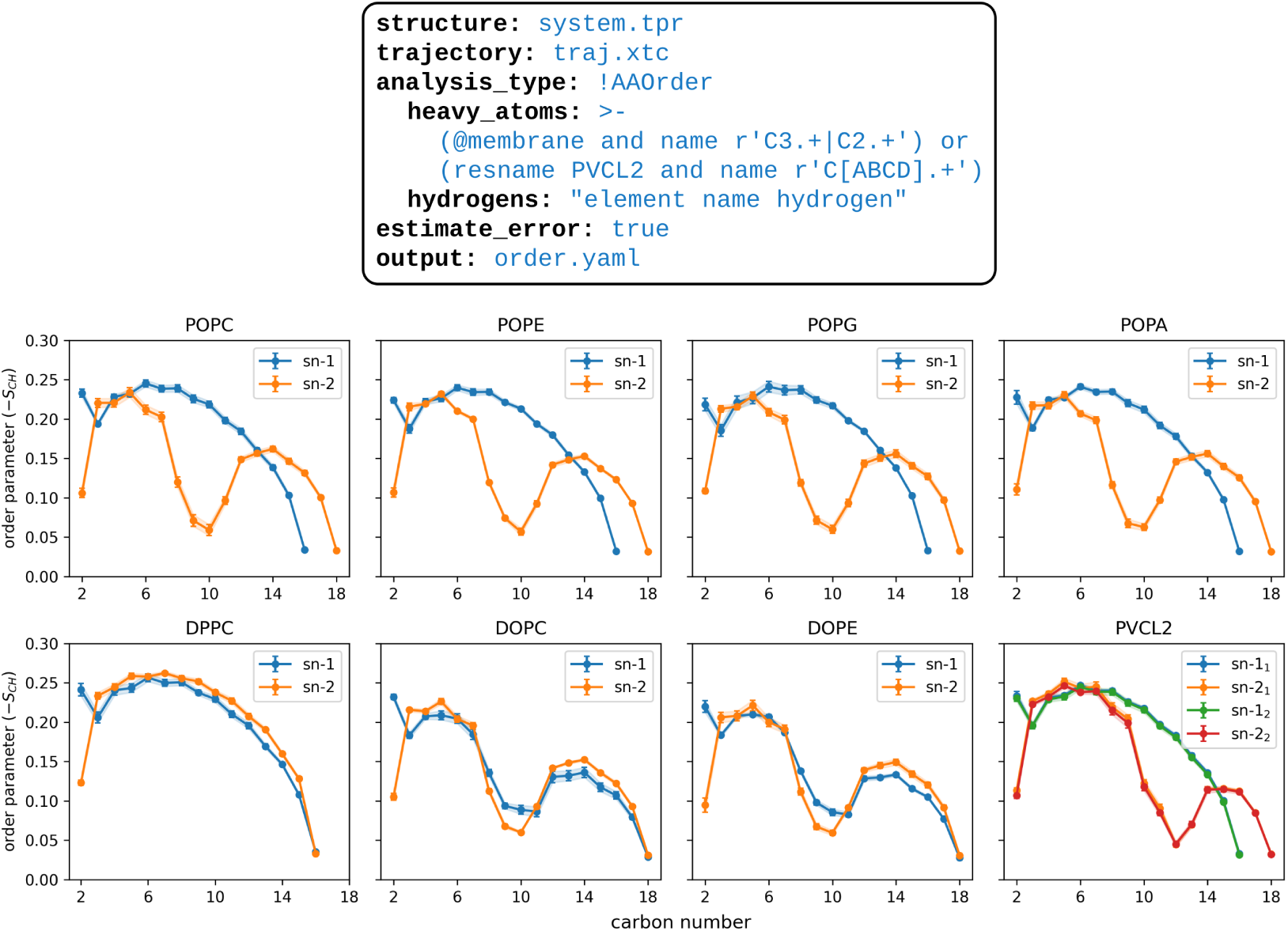
Calculation of order parameters in an atomistic membrane with complex lipid composition. **Top:** Configuration YAML file specifying the analysis parameters for gorder. The query for heavy atoms selects all palmitoyl and oleoyl carbons of all lipid types. The specific lipid types are identified automatically. Error estimation is requested using `estimate error`: true. **Bottom:** Order parameters calculated by `gorder` for carbons of individual lipid molecule types in a single analysis run. Each line corresponds to a specific lipid tail (sn-1, sn-2) of a specific lipid type. Shading represents analysis error estimated by the program using block averaging.

### 3.3. Membrane with a phospholipid scramblase

In this example, we showcase `gorder`’s ability to create two-dimensional order parameter maps that quantify membrane order in individual regions. We also illustrate a simple, flip-flop-safe leaflet classification method. For this demonstration, we use a simulation of a Martini 3 [67] POPC membrane containing a dimer of voltage-dependent anion channel (VDAC), a common outer mitochondrial membrane protein with scramblase activity [73]. Simulation parameters are described in [73]. The analyzed trajectory segment spanned 2.5 µs (25,000 frames). Before analysis, the VDAC dimer in all trajectory frames was centered in the simulation box and RMSD-aligned to its reference structure from the first frame. Without this alignment, the projected order parameters would be spatially averaged out, obscuring the protein’s effects on membrane structure.

See Figure 4 for the configuration YAML file with analysis parameters, calculated order parameters for individual bonds, and examples of xy-plane order parameter projections (“ordermaps”).

**Figure 4:**
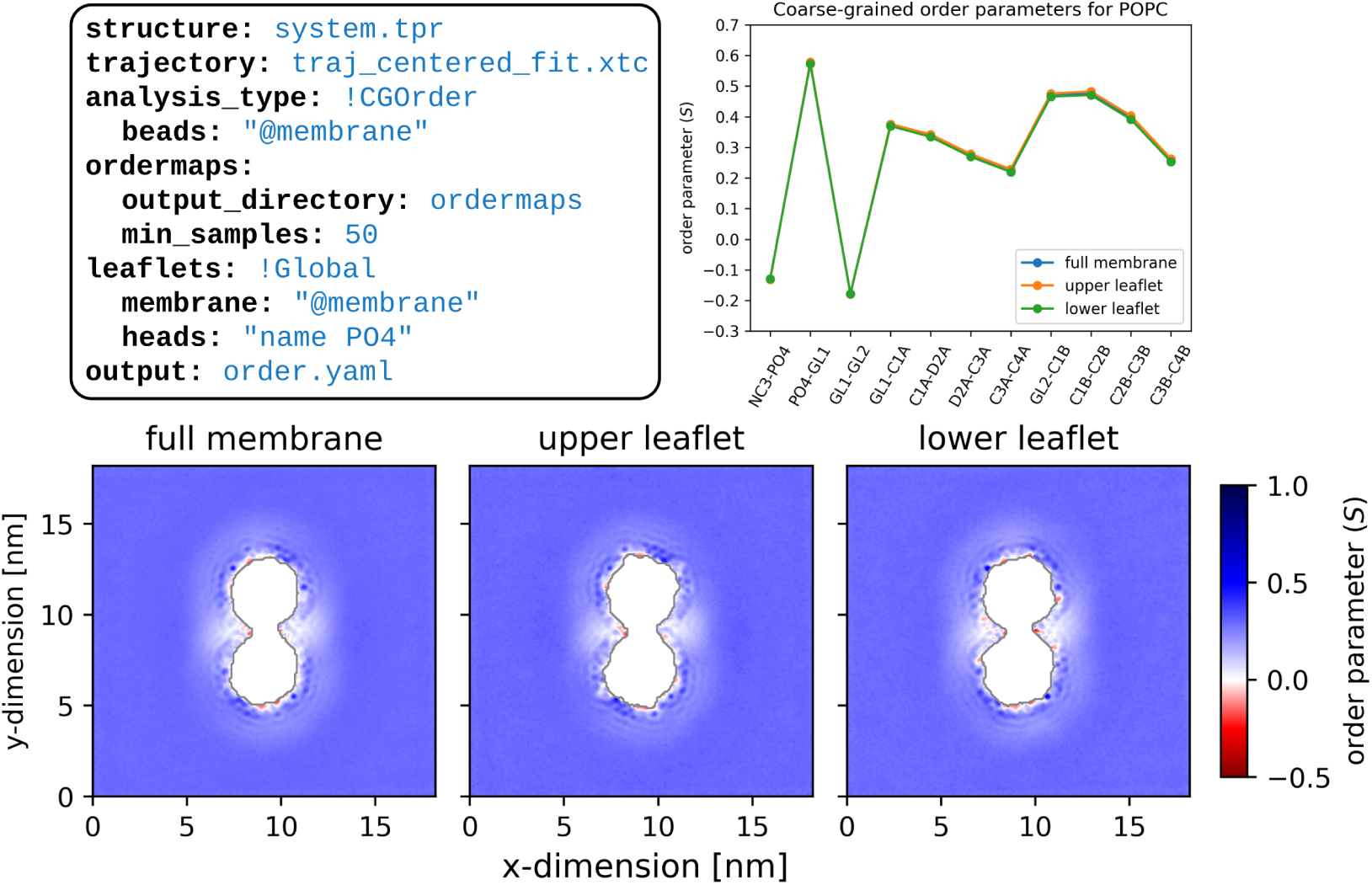
Calculation of order parameters in a coarse-grained membrane containing a phospholipid scramblase. **Top left**: Configuration YAML file specifying the analysis parameters for gorder. The analysis includes all beads of membrane lipids. Order parameter projections (“ordermaps”) are written to the ordermaps directory. Each bin of the “ordermap” requires a minimum of 50 samples; bins with fewer samples are assigned *NaN*. The default bin size is 1×1 Å. Lipid leaflet assignment is determined in each trajectory frame by the position of the selected headgroup atom (name PO4, the phosphate bead) relative to the membrane’s center of geometry. **Top right**: Coarse-grained order parameters for individual POPC lipid bonds in the full membrane, upper leaflet, and lower leaflet. The order parameters are nearly identical between leaflets. **Bottom**: xy-plane projections (“ordermaps”) of average order parameters for all bonds of all lipid molecules in the full membrane and in each leaflet. The gray-outlined white area in each chart represents the protein dimer, where no lipids are present. Protein presence disrupts local membrane structure, visible as reduced order parameters (*S* ≈ 0) or even anti-order (*S <* 0) near the protein. While only average order parameter maps are shown here, gorder additionally generates individual maps for each bond type and for heavy atom types in atomistic/united-atom systems.

### 3.4. Buckled membrane

This example demonstrates gorder’s ability to calculate order parameters in buckled membranes, identify leaflets in such curved systems, and compute order parameters for membrane region within a specified geometric shape. We analyzed a segment of an atomistic simulation of an artificially buckled POPC membrane (1004 lipid molecules, CHARMM36 force field [72], 400 ns duration, 10,000 frames). Prior to analysis, the membrane was aligned to fix the positions of its curvature maximum and minimum. For full simulation parameters, see the Supporting Information.

As shown in Figure 5, the average order parameters in a buckled membrane are nearly identical to those in a flat membrane of the same composition. However, dramatic differences emerge when comparing regions with positive versus negative curvature. Specifically, lipid tails in positively curved regions exhibit significantly higher order than those in negatively curved regions. This is because positive curvature compresses lipid tails relative to their headgroups, enforcing more ordered conformations, whereas negative curvature expands the conformational space available to the tails, resulting in disordering. These results also agree with previous findings [74].

**Figure 5:**
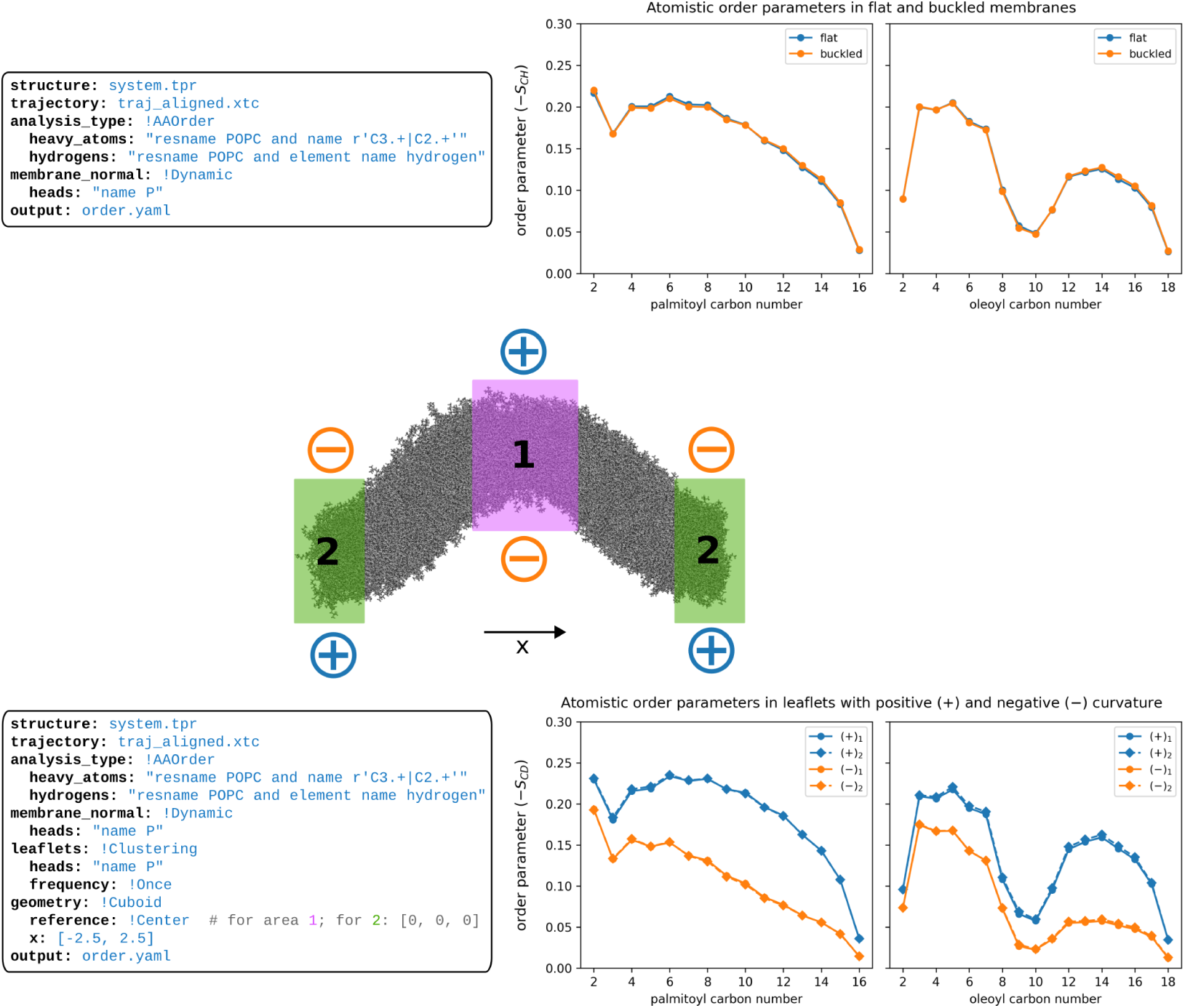
Calculation of order parameters in an atomistic buckled membrane. **Top left:** Configuration YAML file specifying analysis parameters, performing calculations with dynamic membrane normals (no leaflet assignment). All lipids in the membrane are analyzed, no matter their position. **Top right:** Order parameter comparison between buckled and flat POPC membranes. Average values remain nearly identical despite curvature differences. **Center:** Simulation snapshot showing the buckled membrane with labeled leaflet curvatures ((+) convex, (−) concave) and regions analyzed in the next part (purple and green rectangles). **Bottom left:** Configuration YAML file for region-specific analysis. Two analyses were performed, selecting bonds within one of two cuboidal regions. Both regions extend infinitely in the yz-plane while spanning 5 nm along the x-axis and are centered at either the simulation box center (region 1, purple) or the box origin (region 2, green). Lipid leaflet assignment uses the clustering method, performed only for the first frame (due to absent lipid flip-flop) and applied to all subsequent frames. **Bottom right:** Curvature-dependent order parameters. Identical leaflet curvatures yield matching order parameters in both regions, while opposing curvatures show significant differences, as discussed in the main text.

`gorder` can also calculate order parameters for membrane vesicles, as shown in Figure S8. Here, the trends are similar to the buckled membrane, with the inner (negatively curved) leaflet displaying reduced order parameters compared to the outer leaflet.

## 4. Impact

Lipid order parameters are commonly calculated properties of membrane systems, typically used for force field development [1, 2, 3, 4, 5, 6, 7, 67], characterization of membrane structural properties [8, 9, 10, 11, 12, 13, 14, 15], or validation of new simulation approaches [18, 19, 20, 30, 38].

With gorder, we bundle together the features of previously existing programs for lipid order calculations available in the GROMACS community while reimplementing them in a computationally efficient way. Our primary goal is to eliminate the problematic usage of the gmx order tool and the artifacts it produces. Although these artifacts may not affect the scientific validity of the results in most cases, their frequent neglect is concerning. The continued widespread use of gmx order indicates a clear need for a lipid order analysis tool that is more robust, while remaining at least equally simple to use and at least as computationally efficient. We believe that gorder fulfills these requirements completely.

Moreover, gorder’s features enable comprehensive exploration of membrane properties across diverse systems. By automatically identifying lipid types and calculating order parameters separately for each of them in a single pass through the trajectory, gorder streamlines the analysis of membranes with complex lipid compositions—systems that are increasingly common in molecular simulations. Combined with its computational efficiency, gorder is particularly well-suited for analyzing large, complex membranes with biologically relevant lipid compositions.

The ability of gorder to dynamically identify membrane leaflets, even in curved membranes, also simplifies the characterization of structural properties of individual leaflets, especially in membranes with lipid flip-flop. Furthermore, the tool facilitates the characterization of membrane properties near transmembrane proteins or other structures through two-dimensional order parameter projections and by enabling calculations in dynamically selected membrane regions (e.g., in the vicinity of the protein). Notably, gorder can calculate lipid order parameters in membranes with curved geometries, such as vesicles, buckled membranes, or tubes, enabling robust structural characterization of lipids in these complex systems.

We believe gorder is useful even for basic analysis tasks—arguably already relatively well-supported by other simulation programs—as it offers unprecedented speed and various quality-of-life improvements. These include automatic trajectory concatenation, seamless handling of periodic boundary conditions in orthogonal simulation boxes, and flexible selection of time range for analysis. gorder is independent of force field and resolution eliminating the need for multiple specialized programs for different simulation approaches. Finally, gorder demonstrates that the Rust programming language can effectively support the development of complex, high-performance scientific software for analysis of molecular simulations—a domain traditionally dominated by C, C++, and Fortran.

## 5. Conclusions

In this work, we introduce gorder, a new computational tool for calculating lipid order parameters in atomistic, united-atom, and coarse-grained molecular systems, with primary support for GROMACS simulations.

The gorder tool is completely force field independent, has superior computational performance compared to existing alternatives, and provides extensive functionality for performing complex analyses of diverse membrane systems, including those with non-planar geometries.

gorder is distributed as free software under the MIT License. It is available in multiple forms: as a command-line application, a Python package, and a Rust crate. The tool can be obtained from crates.io/crates/gorder and github.com/Ladme/gorder, with comprehensive documentation available at ladme.github.io/gorder-manual. The offline version of the manual and raw results from the performed analyses are available from doi.org/10.5281/zenodo.15282375.

## CRediT authorship contribution statement

**Ladislav Bartš:** Conceptualization; Methodology; Investigation; Data curation; Software; Writing – original draft preparation. **Peter Pajtinka:** Methodology; Investigation; Validation; Writing – original draft preparation; Writing – review & editing. **Robert Vácha:** Funding acquisition; Project administration; Resources; Supervision; Writing – review & editing.

## Declaration of Generative AI and AI-assisted technologies in the writing process

During the preparation of this work, the authors used large language models for editorial corrections and proofreading of the manuscript. The authors reviewed and edited the generated content and take full responsibility for it.

## Acknowledgements

We thank the members of the RoVa Research Group for testing and feature suggestions. The work was supported by the European Research Council (ERC) under the European Union’s Horizon 2020 research and innovation programme (Grant Agreement No. 101001470) and the project National Institute of virology and bacteriology (Programme EXCELES, ID Project No. LX22NPO5103) – Funded by the European Union – Next Generation EU. Computational resources were provided by the CESNET, CERIT Scientific Cloud, and IT4 Innovations National Supercomputing Center by MEYS CR through the e-INFRA CZ (ID: 90254).

