## Supporting Information for "gorder: Comprehensive tool for calculating lipid order parameters from molecular simulations"

### S1. Selected computational approaches and algorithms

This section details selected algorithms and computational methods implemented in `gorder`, including hydrogen position predictions, leaflet classification, dynamic membrane normal calculation, and parallel processing via multithreading.

#### S1.1. Predicting hydrogen positions

`gorder` uses the same geometric reconstruction approach as `buildH` [1] to predict hydrogen positions in united-atom systems. Below, we outline the methodology for all supported groups. Note that all vectors specified in the description have unit lengths. For further details, see the `buildH` documentation at buildh.readthedocs.io.

##### Methyl ($CH_3$) group

Hydrogen reconstruction for $CH_3$ requires two helper atoms: helper1 (a heavy atom directly bonded to the target carbon, C) and helper2 (a heavy atom bonded to helper1). The rotation axis is defined as $\mathbf{v}_{\text{rot}} = (\text{C} \to \text{helper2}) \times (\text{C} \to \text{helper1})$. The first hydrogen is positioned 1.09 Å from C in the direction obtained by rotating C → helper1 by the tetrahedral angle $\arccos(-1/3)$ ($\sim$109.47°) about $\mathbf{v}_{\text{rot}}$. The remaining two hydrogens are generated by rotating this position by $\pm\frac{2\pi}{3}$ radians about C → helper1, preserving tetrahedral geometry and bond lengths.

##### Methylene ($CH_2$) group

For $CH_2$, helper1 and helper2 (heavy atoms bonded to C) determine the reconstruction. The plane normal vector is $\mathbf{v}_{\text{normal}} = (\text{C} \to \text{helper2}) \times (\text{C} \to \text{helper1})$, and the rotation axis is $\mathbf{v}_{\text{axis}} = (\text{C} \to \text{helper1}) - (\text{C} \to \text{helper2})$. The vector $\mathbf{v}_{\text{rotate}} = \mathbf{v}_{\text{normal}} \times \mathbf{v}_{\text{axis}}$ is rotated by $\pm\frac{\theta}{2}$ ($\theta = \arccos(-1/3)$) about $\mathbf{v}_{\text{axis}}$ to position hydrogens 1.09 Å from C, enforcing tetrahedral geometry.

##### Methanetriyl (CH) group

For tertiary CH, three helper atoms (helper1, helper2, helper3) bonded to C determine the hydrogen direction. The resultant vector $\mathbf{v}_{\text{CH}} = -\sum_{X=1}^{3}(\text{C} \to \text{helperX})$ defines the C–H bond. The hydrogen is placed 1.09 Å away from C along $\mathbf{v}_{\text{CH}}$.

##### Methine (unsaturated CH) group

For unsaturated CH, helper1 and helper2 (carbon atoms bonded to the target carbon, C) define the angle $\gamma = \angle$(helper1-C-helper2). The rotation axis $\mathbf{v}_{\text{axis}} = (\text{C} \rightarrow \text{helper1}) \times (\text{C} \rightarrow \text{helper2})$ is used to rotate C $\rightarrow$ helper2 by $\pi - \frac{\gamma}{2}$ radians, bisecting $\gamma$. The hydrogen is positioned 1.09 Å from C along this bisector.

##### Note on helper atom ordering

The selection and sequence of helper atoms is automatically determined based on their sequence in the input topology. For $CH_3$ groups, this can lead to different order parameters for individual C–H bonds calculated by `gorder` compared to `buildH`, where the selection and sequence of helper atoms must be explicitly provided. However, united-atom order parameters for individual C-H bonds of methyl groups inherently lack reliability anyway, as noted in the `gorder` manual. Crucially, the average order parameter for the methyl carbon remains unaffected by helper atom selection.
For $CH_2$ groups, `gorder` may return pro-R/pro-S hydrogen assignments in reversed order relative to `buildH`. This reversal can occur inconsistently across carbons within the same molecule, depending on topology-specific helper atom sequences. While this affects individual bond labels, the calculated order parameters remain valid – only their correspondence to specific predicted hydrogens becomes dependent on specific topology definition.
The sequence of helper atoms has no effect on the predictions in the saturated and unsaturated CH groups.

#### S1.2. Leaflet classification methods

When requested, `gorder` dynamically identifies membrane leaflets and performs separate analyses for each leaflet using one of four classification methods: "global", "local", "individual", or "clustering".

##### Global method

The global method requires the user to provide a selection of all membrane atoms and a selection of lipid headgroup atoms (one per lipid). Lipids are assigned to the upper leaflet if the shortest oriented distance between any periodic image of their headgroup and the membrane center of geometry (along the membrane normal direction) is positive; otherwise, they are assigned to the lower leaflet. The membrane center of geometry is calculated using the refined Bai and Breen algorithm [2, 3], which provides accurate results in periodic systems even when the atom selection spans periodic boundaries. This same algorithm is used throughout `gorder` whenever a center of geometry calculation is needed, such as when selecting membrane regions and using group centers as reference points. The global method is recommended for planar membranes with undulations smaller than the membrane thickness.

##### Local method

The local method requires the user to provide a selection of all membrane atoms, a selection of lipid headgroup atoms (one per lipid), and a cylinder radius. Lipid assignment follows the same principle as the global method, but uses a locally calculated membrane center of geometry. This local center is determined using membrane atoms within a cylinder centered at the headgroup atom, aligned with the membrane normal, and having the user-specified radius and an infinite height. Cell lists are used to optimize the atom selection process, reducing the theoretical time complexity from $\mathcal{O}(n^2)$ to $\mathcal{O}(n)$. Despite this optimization, the local method is much slower than both the global and the individual method, and should only be used for planar membranes where those methods fail.

##### Individual method

The individual method requires selections for lipid headgroup atoms (one per lipid) and tail-end atoms representing a terminal segment of each lipid tail (any number per lipid). For each lipid molecule, the method sums the shortest oriented distances (along the membrane normal) between the headgroup atom (or its periodic images) and all tail-end atoms along the membrane normal. A positive total distance assigns the lipid to the upper leaflet; a negative distance assigns it to the lower leaflet. The individual method is very fast and is recommended for large planar or undulating membranes.

##### Clustering method

The clustering method uses only a selection of lipid headgroup atoms, applying spectral clustering [4] to divide them (and the corresponding lipid molecules) into two clusters (leaflets). Headgroup atoms are treated as graph nodes, with edge weights inversely proportional to interatomic distances. After constructing the Laplacian matrix from these weights, its two smallest non-zero eigenvectors are identified (via either full eigendecomposition or the approximate Lanczos method [5, 6]) and clustered using k-means clustering. Cluster with the higher number of lipid molecules or containing the lowest atom index in the first analyzed trajectory frame is considered to be the upper leaflet.

Clustering method is the only method implemented in `gorder` able to identify leaflets in curved membranes. Unlike MDAnalysis's LeafletFinder [7], the clustering method can handle membranes with flip-flopping lipids and

lipid pores but carries higher computational cost. `gorder` attempts to mitigate this by using cell lists to sparsify the Laplacian (considering only nearby atom pairs), and automatically selecting between full and approximate eigendecomposition based on system size and precision requirements.
Note that implementation details—including parameters for the calculation of edge weights, distance cutoffs for selecting nearby lipids, and the eigenvector computation strategy—may change between `gorder` versions.

#### S1.3. Dynamic estimation of membrane normals

If requested, `gorder` can dynamically estimate membrane normals for each lipid molecule by selecting lipid headgroups near the molecule's own headgroup (within 2 nm by default) and performing principal component analysis on their positions. The last principal component (the direction of least positional variation) is then assigned as the membrane normal for the selected lipid. The search for nearby lipid headgroups is optimized using cell lists.

#### S1.4. Multithreading

`gorder` utilizes multithreading through Rust's native threads to accelerate the analysis. Although the basic algorithm of order parameters calculation is embarrassingly parallelizable since the trajectory can be divided into $N$ blocks that are read and analyzed in parallel, the number of frames in a trajectory is unknown until processed, complicating problem decomposition. To avoid having to read the trajectory multiple times, `gorder` employs a single-pass strategy with interleaved frame allocation: threads divide the workload by processing every $N$-th consecutive frame, where $N$ is the number of threads (e.g., thread 1: frames 1, $N$+1, 2$N$+1. . . ; thread 2: frames 2, $N$+2, 2$N$+2. . . ). This requires efficient frame skipping, enabled by the `groan_rs` library. The interleaved approach also allows thread synchronization when performing leaflet assignment for lipids at user-defined intervals (e.g., every 10 frames). See Figure S1 for a depiction of the multithreading scheme employed by `gorder`.

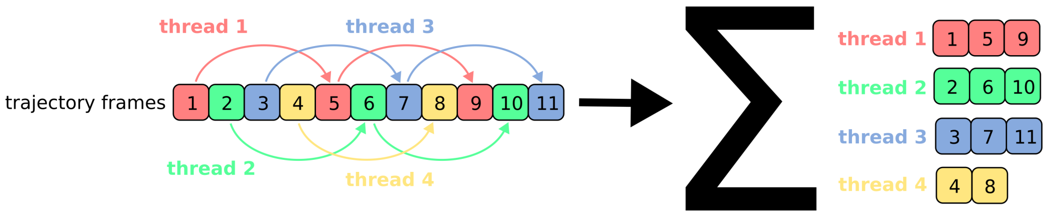


Figure S1: Schematic depiction of the multithreading scheme employed by `gorder`. Frames are allocated to individual threads in an interleaved manner. Each thread then performs the analysis independently, skipping to the frames it is assigned to read. Finally, data from all threads are combined, and the final result is reported.

### S2. Additional examples, validation, and benchmarks

#### S2.1. Order parameters for individual carbon-hydrogen bonds

As noted in the main text, `gorder` reports order parameters for both individual heavy atom types (typically carbons) and specific carbon-hydrogen bond types. This capability is shared by NMR Lipids [8] tools (e.g., the `calc_order` script). Here, we compare order parameters calculated for individual bond types by `gorder` and `calc_order` using the system and trajectory described in Section 3.1.

Figure S2 demonstrates perfect agreement between the order parameters calculated by `gorder` and NMR Lipids' `calc_order` for individual bond types.

#### S2.2. Single-component united-atom membrane

In this example, we analyzed a united-atom POPC membrane (256 lipids, Berger force field, roughly 44,300 atoms) from the NMR Lipids DataBank (zenodo.org/records/1402417). The 300 ns trajectory comprised 3,000 frames. Figure S3 shows the configuration YAML file used for the analysis by `gorder`, analysis results compared with other software, and execution time benchmarks.

#### S2.3. Single-component coarse-grained membrane

In this example, we analyzed a coarse-grained POPC membrane (512 lipids, Martini 3 force field [9], roughly 16,800 beads). The 1 µs trajectory comprised 10,000 frames. See Section S5.2 for complete simulation parameters. Figure

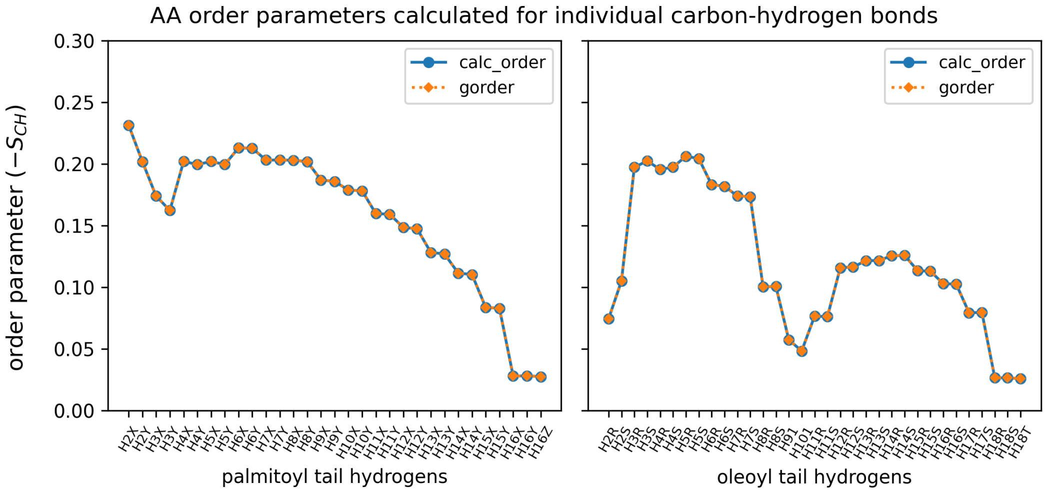


Figure S2: Comparison of order parameters for individual carbon-hydrogen bond types of POPC calculated using `gorder` and NMR Lipids' `calc_order` script. Order parameters calculated by both tools are identical.

S4 shows the configuration YAML file used for the analysis by `gorder`, analysis results compared with other software, and execution time benchmarks.

#### S2.4. United-atom order parameters in an atomistic membrane

In this example, we perform additional validation of united-atom order parameter calculations by direct comparison with atomistic order parameters calculated from the same simulation (using the system described in Section 3.1). Figure S5 displays the configuration YAML file for united-atom parameter calculations using a simulation of an atomistic CHARMM36 [10] membrane, and a comparison of calculated united-atom and atomistic order parameters.

#### S2.5. Performance with increasing membrane complexity

As mentioned in the main text, `gorder` calculates order parameters for all lipid types in a membrane simultaneously during a single trajectory pass. Here we evaluate `gorder`'s performance with increasing membrane complexity (i.e., number of distinct lipid types). We compare single-threaded (1 thread) and parallel (8 threads) executions across three simulations (200

```yaml
structure: system.tpr
trajectory: traj.xtc
analysis_type: !UAOrder
  saturated: >-
    resname POPC and name C17 C18 C19 C20 C21 C22
    C23 C26 C27 C28 C29 C30 C31 CA1 CA2 C36 C37 C38
    C39 C40 C41 C42 C43 C44 C45 C46 C47 C48 C49 C50
  unsaturated: "resname POPC and name C24 C25"
output: order.yaml
```

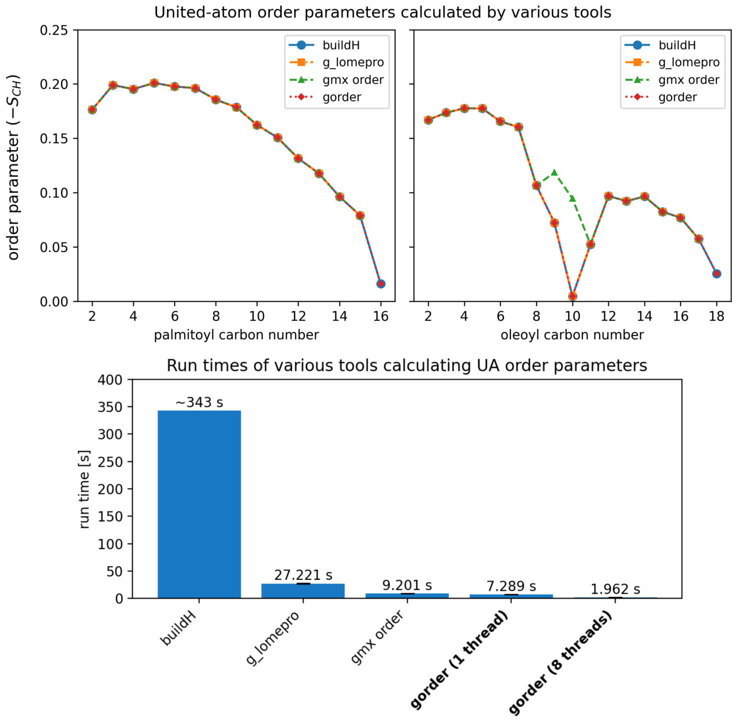


Figure S3: Calculation of order parameters in a united-atom (Berger lipids) system using `gorder` and comparison with other software. **Top:** Configuration YAML file specifying the analysis parameters for `gorder`. The query for `saturated` selects saturated carbons of palmitoyl and oleoyl tails, while the query for `unsaturated` selects unsaturated carbons. **Center:** Order parameters calculated by `gorder` for palmitoyl and oleoyl tails of POPC (red line), compared with results from other tools (`buildH` script, `g_lomepro` program, and `gmx order`). All tools except `gmx order` yield consistent results. `gmx order` should only be used for fully saturated tails of united-atom lipids. **Bottom:** Execution times for order parameter calculations of POPC tail carbons across different tools. Benchmarking was performed on GNU/Linux Mint 20.2 (8-core Intel Core i7-11700, Samsung 870 EVO SSD) with cold cache using `hyperfine`. `gorder` was compiled using `rustc` v1.87.0 and `gcc` v9.4.0. `gorder`, `gmx order`, and `g_lomepro` were executed 5 times each, while `buildH` was much slower and was run once. Note that `g_lomepro` and `gmx order` require two runs to analyze both lipid tails. For single-tail analysis (single run), `gorder` is marginally faster than `gmx order` (`gorder`: 4.02 s, `gmx order`: 4.75 s). Note that unlike `gmx order`, `gorder` also reports order parameters for individual predicted carbon-hydrogen bonds.

ns, 10,000 frames each) of CHARMM36 [10] membranes with varying com-

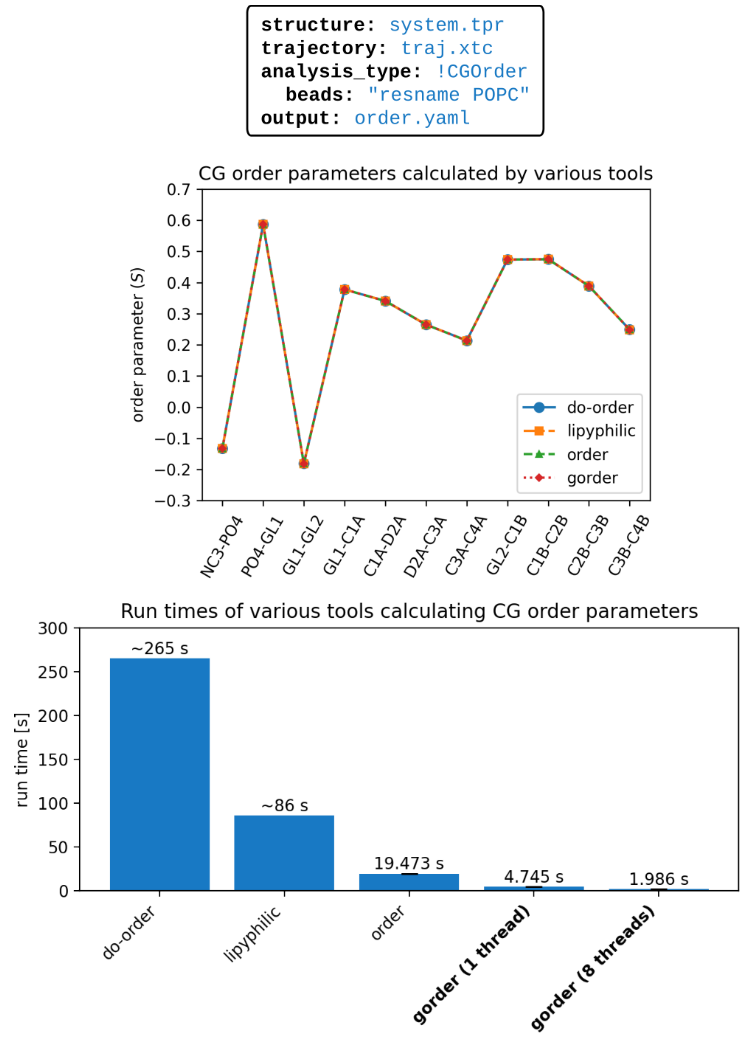


Figure S4: Calculation of order parameters in a coarse-grained (Martini 3) system using `gorder` and comparison with other software. **Top:** Configuration YAML file specifying the analysis parameters for `gorder`. The query for `beads` selects all beads of common membrane lipids. **Center:** Order parameters calculated by `gorder` for all bonds of POPC lipids (red line), compared with results from other tools (Martini's `do-order` script, `lipyphilic` library, and `order` program). All tools yield consistent results. **Bottom:** Execution times for calculations of coarse-grained order parameters for POPC bonds across different tools. Benchmarking was performed on GNU/Linux Mint 20.2 (8-core Intel Core i7-11700, Samsung 870 EVO SSD) with cold cache using `hyperfine`. `gorder` was compiled using `rustc` v1.87.0 and `gcc` v9.4.0. `gorder` and `order` were executed 5 times each, while the other two, slower tools were run once. Note that the `lipyphilic` library requires 11 runs to analyze all bonds of POPC lipids, as it only reports average order parameters for all selected bonds irrespective of their type. Even when only average order parameters are needed, `gorder` remains faster.

```yaml
structure: system.tpr
trajectory: traj.xtc
analysis_type: !UAOrder
  saturated: >-
    resname POPC and name r'C3.+|C2.+' and
    not name C21 C31 C29 C210
  unsaturated: "resname POPC and name C29 C210"
  ignore: "element name hydrogen"
output: order.yaml
```

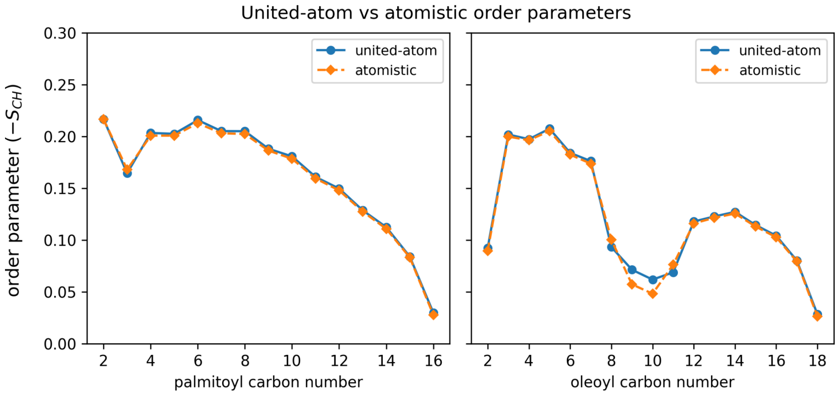


Figure S5: Calculation of united-atom order parameters for an atomistic membrane. **Top:** Configuration YAML file for united-atom order parameter calculation. The `ignore` query excludes all explicit hydrogen atoms from the analysis. **Bottom:** Comparison of united-atom and atomistic order parameters calculated for the same system. Hydrogen position prediction shows high reliability for saturated carbons but lower accuracy for unsaturated carbons. Note that when hydrogen positions are available, atomistic order parameters should *always* be preferred.

plexity: single-component POPC (used in Section 3.1), three-component POPC:POPE:POPG (128:64:64) membrane, and an eight-component system (used in Section 3.2). All systems contained identical lipid counts (256 total) and similar atom numbers (64,500 (single-component) vs. 64,100 (three-component) vs. 67,300 (eight-component)).

Figure S6 reveals no systematic dependence of run time on membrane complexity. While the eight-component membrane analysis is slightly slower, the analysis of the ternary mixture (POPC:POPE:POPG) runs as fast as for the single-component system. We attribute the decreased analysis speed for the eight-component system to the slightly larger number of atoms in it and to the presence of cardiolipin (see Figure S7). Even in the worst case scenario, `gorder`'s complexity scaling is significantly better than the typical $\mathcal{O}(n)$ dependence of other order parameter analysis tools, where $n$ is the number of lipid types.

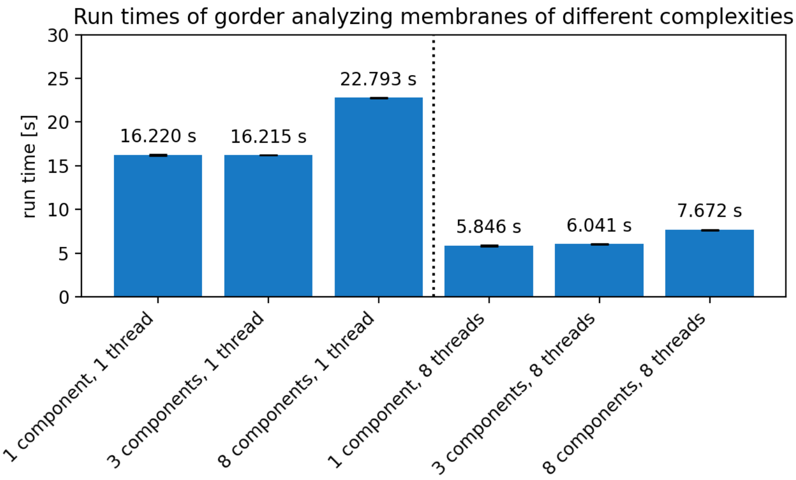


Figure S6: Execution time benchmarking of `gorder` for single-component (POPC), three-component (POPC:POPE:POPG), and eight-component membranes using 1 and 8 threads. The analysis time shows minimal dependence on membrane compositional complexity (number of distinct lipid types). Benchmarking was performed on GNU/Linux Mint 20.2 (8-core Intel Core i7-11700, Samsung 870 EVO SSD) with cold cache using `hyperfine`. `gorder` was compiled using `rustc` v1.87.0 and `gcc` v9.4.0. Each analysis was repeated five times; reported values represent the mean execution time.

#### S2.6. Performance with increasing lipid size

Here, we compare the performance of `gorder` calculating order parameters for membranes composed of lipid types of very different sizes. Specifically, we compare a POPC system (256 lipids, with 64 analyzed C-H bonds in the tails of each lipid) with a cardiolipin (CDL) system (128 lipids, with 128 analyzed C-H bonds in the tails of each lipid). Both systems contained the same number of analyzed bonds (16,384) and similar total numbers of atoms (64,500 atoms in the POPC system versus 61,400 atoms in the CDL system). The analyzed portion of each trajectory was 200 ns long and consisted of 10,000 frames.

Figure S7 shows that the analysis of the cardiolipin-containing system was roughly 25% slower than that of the POPC system, despite both systems having the same number of analyzed bonds and the CDL system containing slightly fewer total atoms.

#### S2.7. Membrane vesicle

In this example, we demonstrate `gorder`'s capability to calculate lipid order parameters in membrane vesicles using a simulated POPC vesicle (Martini

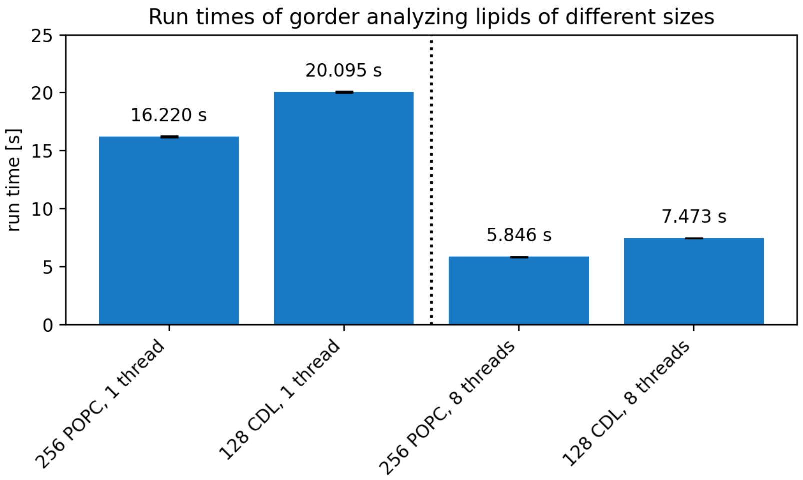


Figure S7: Execution time benchmarking of `gorder` for POPC- and cardiolipin-containing systems using 1 and 8 threads. The same number of bonds was analyzed in each system. The results show that the analysis of cardiolipin, which has twice as many bonds per molecule as POPC, is more than twice as slow as the analysis of the smaller POPC lipid. Benchmarking was performed on GNU/Linux Mint 20.2 (8-core Intel Core i7-11700, Samsung 870 EVO SSD) with cold cache using `hyperfine`. `gorder` was compiled using `rustc` v1.87.0 and `gcc` v9.4.0. Each analysis was repeated five times; reported values represent the mean execution time.

3 force field [9], 2758 lipids, diameter ≈ 20 nm). The analyzed 2 µs trajectory segment comprised 10,000 frames. See Section S5.2 for full simulation parameters.

Figure S8 shows that the outer vesicle leaflet maintains order parameters comparable to a flat membrane of identical composition. In contrast, the inner leaflet exhibits significantly reduced tail order. This disorder arises from the expanded conformational space available to lipid tails in the inner leaflet, mirroring the behavior observed in negatively curved regions of the buckled membrane (see Section 3.4). The presented findings also agree with previously published results [11].

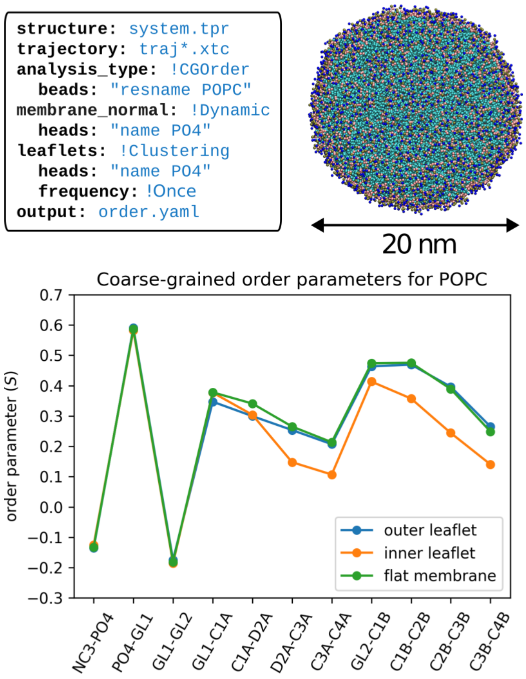


Figure S8: Calculation of order parameters in a coarse-grained membrane vesicle. **Top left:** Configuration YAML file specifying analysis parameters, including dynamic membrane normal calculation and leaflet classification using the clustering method. Note that all trajectory files matching the glob pattern `traj*.xtc` are selected, seamlessly concatenated, and analyzed. **Top right:** Simulation snapshot showing the analyzed vesicle with approximate diameter indicated. **Bottom:** Comparison of order parameters between the outer and inner vesicle leaflets and a planar membrane of identical lipid composition.

### S3. Configuration YAML files used

The YAML configuration files used for all `gorder` analyses in the article are provided below in a copy-ready format. To run the analysis, save the configuration as `analyze.yaml`, and execute the command `gorder analyze.yaml` in your terminal.

Listing 1: Input parameters for Figures 2 and S2

```
structure: system.tpr
trajectory: traj.xtc
analysis_type: !AAOrder
  heavy_atoms: "resname POPC and name r'C3.+|C2.+'"
  hydrogens: "resname POPC and element name hydrogen"
output: order.yaml
# n_threads: 8  # uncomment to perform using 8 threads
```

Listing 2: Input parameters for Figure 3

```
structure: system.tpr
trajectory: traj.xtc
analysis_type: !AAOrder
  heavy_atoms: >-
    (@membrane and name r'C3.+|C2.+') or
    (resname PVCL2 and name r'C[ABCD].+')
  hydrogens: "element name hydrogen"
estimate_error: true
output: order.yaml
```

Listing 3: Input parameters for Figure 4

```
structure: system.tpr
trajectory: traj_centered_fit.xtc
analysis_type: !CGOrder
  beads: "@membrane"
ordermaps:
  output_directory: ordermaps
  min_samples: 50
leaflets: !Global
  membrane: "@membrane"
  heads: "name PO4"
output: order.yaml
```

Listing 4: Input parameters for Figure 5 (no assignment)

```
structure: system.tpr
trajectory: traj_aligned.xtc
analysis_type: !AAOrder
  heavy_atoms: "@membrane and name r'C3.+|C2.+'"
  hydrogens: "@membrane and element name hydrogen"
membrane_normal: !Dynamic
  heads: "name P"
output: order.yaml
```

Listing 5: Input parameters for Figure 5 (area 1/2)

```
structure: system.tpr
trajectory: traj_aligned.xtc
analysis_type: !AAOrder
  heavy_atoms: "@membrane and name r'C3.+|C2.+'"
  hydrogens: "@membrane and element name hydrogen"
membrane_normal: !Dynamic
  heads: "name P"
leaflets: !Clustering
  heads: "name P"
  frequency: !Once
geometry: !Cuboid
  reference: !Center    # for area 1
# reference: [0, 0, 0]  # uncomment for area 2
  x: [-2.5, 2.5]
```

Listing 6: Input parameters for Figure S3

```
structure: system.tpr
trajectory: traj.xtc
analysis_type: !UAOrder
  saturated: >-
    resname POPC and name C17 C18 C19 C20 C21 C22
    C23 C26 C27 C28 C29 C30 C31 CA1 CA2 C36 C37 C38
    C39 C40 C41 C42 C43 C44 C45 C46 C47 C48 C49 C50
  unsaturated: "resname POPC and name C24 C25"
output: order.yaml
# n_threads: 8  # uncomment to perform using 8 threads
```

Listing 7: Input parameters for Figure S4

```
structure: system.tpr
trajectory: traj.xtc
analysis_type: !CGOrder
  beads: "resname POPC"
output: order.yaml
# n_threads: 8  # uncomment to perform using 8 threads
```

Listing 8: Input parameters for Figure S5

```
structure: system.tpr
trajectory: traj.xtc
analysis_type: !UAOrder
  saturated: >-
    resname POPC and name r'C3.+|C2.+' and
    not name C21 C31 C29 C210
  unsaturated: "resname POPC and name C29 C210"
  ignore: "element name hydrogen"
output: order.yaml
```

Listing 9: Input parameters for Figure S8

```
structure: system.tpr
trajectory: traj*.xtc
type: !CGOrder
  beads: "resname POPC"
membrane_normal: !Dynamic
  heads: "name PO4"
leaflets: !Clustering
  heads: "name PO4"
  frequency: !Once
output: order.yaml
```

### S4. Scripts, programs, and libraries used for validation

The results of the analyses performed by `gorder` and the program's speed were compared with those of other tools, as mentioned in the main text and in the previous sections. Below is a list of all these tools and their sources:

- `calc_order` by NMR Lipids available from `github.com/NMRLipids/Databank/blob/6a91be2270e89ec7bb9c75006c2f2a2507c24a01/Scripts/BuildDatabank/OrderParameter.py`; the script is also available from `doi.org/10.5281/zenodo.15282375`
- VMD's `calc_op.tcl` available from `www.ks.uiuc.edu/Research/vmd/mailing_list/vmd-l/att-14731/calc_op.tcl`; also available from `doi.org/10.5281/zenodo.15282375`
- LOOS library's `order_params` available from `github.com/GrossfieldLab/loos/releases/tag/v4.2.0`
- `gmx order` available as part of GROMACS version 2021.4 from `github.com/gromacs/gromacs/releases/tag/v2021.4`
- `buildH` version 1.6.1 available from `github.com/patrickfuchs/buildH/releases/tag/v1.6.1`
- `g_lomepro` package available from `github.com/vgapsys/g_lomepro/tree/5c4bf3817036aa58119d3f43ceb45daef0c0271e`
- `do-order` script available from `cgmartini.nl/docs/downloads/tools/other-tools.html#do-order`, modified to be compatible with the analyzed lipids; also available from `doi.org/10.5281/zenodo.15282375`
- `lipyphilic` library version 0.10 available from `github.com/p-j-smith/lipyphilic/releases/tag/v0.10.0`
- `order` program available from `zenodo.org/records/8369479`

### S5. Molecular dynamics simulations details

#### S5.1. Atomistic simulations

Atomistic simulations were performed using GROMACS 2021.4 (most simulations) and 2020.3 (buckled membrane) [12]. Membranes were represented using the CHARMM36 [10] force field, prepared using CHARMM-GUI Membrane Builder [13, 14, 15] (NaCl concentration of 0.154 mol dm$^{-3}$), minimized using the steepest-descent algorithm, and equilibrated for 7.75 ns in six stages (I-III: 250 ps each, IV-V: 1 ns each, VI: 5 ns) with a slowly decreasing strength of position and dihedral restraints applied to lipid molecules. The first two stages (0.5 ns) were performed in the NVT ensemble, and the next four stages in the NPT ensemble. Each membrane was then simulated for at least 1 µs with production parameters, but only a specified portion of the trajectory was analyzed. The simulation time step was 1 fs in the first three stages of equilibration and 2 fs in all the following stages. The stochastic velocity rescaling thermostat [16] (coupling constant 0.5 ps, temperature 310 K, two coupling groups: membrane and water with ions) was used in all equilibration stages and in the production stage of the simulations. The Berendsen barostat [17] (semi-isotropic coupling scheme ($xy$, $z$), coupling constant: 5 ps, compressibility: $4.5 \times 10^{-5}$ bar$^{-1}$, reference pressure: 1 bar) was used in the equilibration NPT stages. The Parrinello-Rahman barostat [18, 19] (same parameters as Berendsen) was used in the production stage. Bonds with hydrogens were constrained using LINCS [20].

##### Used lipids

The following lipid types were used in the simulated membranes:

- 1-palmitoyl-2-oleoyl-*sn*-glycero-3-phosphocholine (POPC)
- 1-palmitoyl-2-oleoyl-*sn*-glycero-3-phosphoethanolamine (POPE)
- 1-palmitoyl-2-oleoyl-*sn*-glycero-3-phosphoglycerol (POPG)
- 1-palmitoyl-2-oleoyl-*sn*-glycero-3-phosphate (POPA)
- 1,2-dioleoyl-*sn*-glycero-3-phosphocholine (DOPC)
- 1,2-dioleoyl-*sn*-glycero-3-phosphoethanolamine (DOPE)
- 1,2-dipalmitoyl-*sn*-glycero-3-phosphocholine (DPPC)
- 1,3-bis(1-palmitoyl-2-oleoyl-*sn*-glycero-3-phospho)-glycerol (PVCL2)

**Preparation of buckled membrane**

First, a coarse-grained flat membrane patch comprising 1004 POPC molecules was generated using CHARMM-GUI Martini Maker [13, 21]. Martini non-polarizable water beads and ions were then added to achieve an ion concentration of 0.15 mol dm$^{-3}$. The system was energy-minimized using the steepest-descent algorithm. The initially square membrane patch was transformed into a rectangular shape by simultaneously compressing along the y-direction and expanding along the x-direction using the PLUMED plugin (version 2.3) [22] while preserving the membrane area.

Subsequently, the membrane was compressed along the x-direction using PLUMED, with the y-dimension kept fixed, to induce buckling and achieve a compressional strain of 0.16. The buckled membrane was then equilibrated for 200 ns and then simulated for 20 µs. Detailed parameters of these coarse-grained simulations are described in [23].

The buckled membrane system was backmapped to an atomistic representation (CHARMM36) using the CHARMM-GUI All-atom Converter [13]. The system was equilibrated for 1.875 ns in six stages (I-III: 125 ps each; IV-VI: 500 ps each), with progressively decreasing strength of position and dihedral restraints applied to the lipid molecules. The simulation parameters used in these stages correspond to those described above.

Equilibration was followed by a production run. Most simulation parameters remained identical to those used for other atomistic systems, except that the compressibility in the xy-plane was set to zero to maintain the buckled membrane shape. The system was simulated for over 6.5 µs, with only the last 400 ns being analyzed here.

#### S5.2. Coarse-grained simulations

Coarse-grained simulations were performed using GROMACS 2021.4 [12] and molecules were represented using the Martini 3 [9] force field.

The analyzed planar POPC membrane was prepared using the `insane` script (available from github.com/Tsjerk/Insane) with an ion concentration of 0.154 mol dm$^{-3}$, minimized using the steepest-descent algorithm, and equilibrated for 32.75 ns in four stages (simulation lengths of 0.5, 1.25, 1, and 30 ns, respectively) with increasing simulation time step (2, 5, 10, and 20 fs, respectively). All stages of equilibration were performed in the NPT ensemble. The membrane was then simulated for 5 µs with production parameters but only the specified portion of the trajectory was analyzed.

The POPC membrane vesicle was prepared using the CHARMM-GUI Martini Vesicle Builder [13, 21] with an ion concentration of 0.154 mol dm$^{-3}$, minimized using the steepest-descent algorithm, and equilibrated for 110 ns in five stages (simulation length of stage I: 30 ns; simulation length of other stages: 20 ns) with increasing simulation time step (2, 5, 10, 20, and 20 fs, respectively). During the equilibration, six pores were maintained using cylindrical inverted flat-bottom restraints with slowly decreasing force constant applied to the tails of all lipids, allowing water equilibration between the inside and the outside of the vesicle and exchange of lipids between leaflets of the vesicle.

For temperature coupling in both the planar membrane and the vesicle and in all stages, the stochastic velocity rescaling thermostat [16] was used (coupling constant: 1 ps, temperature: 310 K, two coupling groups: membrane and water with ions). For pressure coupling in all stages, the stochastic cell rescaling thermostat [24] was used (semi-isotropic coupling scheme ($xy$, $z$) for the planar membrane, isotropic coupling scheme for the vesicle, coupling constant: 12 ps, compressibility: $3 \times 10^{-5}$ bar$^{-1}$, reference pressure: 1 bar). The `lincs-order` and `lincs-iter` parameters were set to 8 and 2 [25], respectively, and the neighbor list parameters were set according to recommendations in [26].

Parameters for the membrane simulation with a phospholipid scramblase are described in [27].
